# Dual Probe Ligation In Situ Hybridization with Rolling-Circle Amplification for High-Plex Spatial Transcriptomics

**DOI:** 10.1101/2025.01.22.634417

**Authors:** Sarah E. Maguire, Joel Credle, Elizabeth M.W. Bertelson, Sunny Lee, Boyoung Cha, Dan Xie, Greg Kirk, Debjit Ray, Logan George, Aditya Suru, Alexandre Maalouf, Chiseko Ikenaga, Thomas Lloyd, Nicolas J. Llosa, H. Benjamin Larman

## Abstract

New biological insights are increasingly dependent upon a deeper understanding of tissue architectures. Critical to such studies are spatial transcriptomics technologies, especially those amenable to analysis of the most widely available human tissue type, formalin-fixed and paraffin-embedded (FFPE) clinical specimens. Here we build on our previous oligonucleotide probe ligation-based approach to accurately analyze FFPE mRNA, which suffers from variable levels of degradation. Ligation *In Situ* Hybridization followed by rolling circle amplification (LISH-Lock’n’Roll or LISH-LnR), provides a streamlined method to detect the spatial location of specific mRNA isoforms within FFPE tissue architectures. Iterative fluorescent probe hybridization and imaging enables highly multiplexed spatial transcriptomic studies, as demonstrated herein for fixed specimens from inclusion body myositis patients and pediatric rhabdomyosarcoma patients. We additionally demonstrate a system of molecular rheostats that can be used to fine tune the performance of the LISH-LnR assay. Combined with LISH-seq and LISH-QC, the LISH-LnR methodology provides a powerful approach to spatial transcriptomics.

## INTRODUCTION

Spatial biology seeks to elucidate the three dimensional organization of cells and their associated biomolecules within the tissue context (*1*). Technical advances in spatial biology have begun to shed light on the role of different cell types and states within complex tissue architectures in health and diseases (*2–4*). Owing to the digital nature of mRNA, highly multiplexed gene expression analysis technologies have so far tended to provide the greatest density of information. Methods that enable the multiplexed visualization of mRNA *in situ* can therefore unlock previously inaccessible biological insights.

Clinical tissue specimens are most often preserved using formalin fixation and paraffin embedding (FFPE). While this process retains tissue architecture and simplifies long-term storage, it is not optimal for preservation of relatively labile mRNA molecules. FFPE mRNA tends to suffer from highly variable amounts of degradation via covalent modifications and strand fragmentation.(*5*) We have previously described Ligation In Situ Hybridization (LISH) for non-spatially resolved sequencing-based or qPCR-based analysis of FFPE mRNA.(*6*) LISH involves the ligation of short DNA probe pairs designed to anneal adjacent and contiguous sequences on target mRNA molecules, for subsequent ligation, release, amplification and analysis. Incorporating ribonucleotides into the 3’-terminus of the acceptor probes permits high efficiency ligation using the T4 RNA Ligase 2 (Rnl2) enzyme (*7*).

Here we demonstrate the technical performance and utility of the LISH-Lock’n’Roll (LISH-LnR) assay using a combined microfluidic, imaging and analysis instrument optimized for high-plex spatial omics studies. Integration of an automated microfluidics and imaging system enables the robust sequential addition and removal of combinatorial probe pools, which occur before and after each round of imaging, respectively. After circularization of LISH probe sets carrying up to four binding sites for fluorescently tagged oligonucleotides, rolling-circle amplification creates many copies of these binding sites *in situ* as a high-fidelity proxy for the original mRNA target. Key features of the rolling circle products (RCPs), including their size and number per target molecule, may also be tuned for specific use cases and instrumentation. Combinatorial detection with fluorescently tagged oligonucleotide readout probes enables specific detection of hundreds to thousands of distinct mRNA targets.

The non-spatially resolved LISH assay can be employed to enhance probe panel design and to pre-qualify the sample mRNA integrity and availability to ensure a successful spatial transcriptomic analysis. These features of the LISH platform are integrated to explore a key splicing event in FFPE resections obtained from pediatric rhabdomyosarcoma patients.

## RESULTS

### Creation and detection of LISH-LnR products *in situ*

Previously, we introduced a LISH probe set design comprised of a (“5P”) phosphorylated “donor” probe and a (“3P”) diribonucleotide-terminated “acceptor” probe.(*6*) Each probe contains a target recognition sequence, designed to hybridize adjacent sequences on the target RNA, and universal appended adapter sequences, used for amplification and downstream sequencing or qPCR. For adaptation to spatially-resolved *in situ* analyses, we modified the LISH-seq probe design by inserting readout probe (RP) binding sequences between the target recognition sequences and the universal adapter sequences (**Fig. 1a**). In the new LISH-Lock’n’Roll (LISH-LnR) assay, the universal adapter sequences are used for probe set circularization (“5P-Bridge” and “3P-Bridge” sequences). Circularization by enzymatic ligation requires the 5’-terminus of the 3P probe to be phosphorylated.

**Figure 1.**
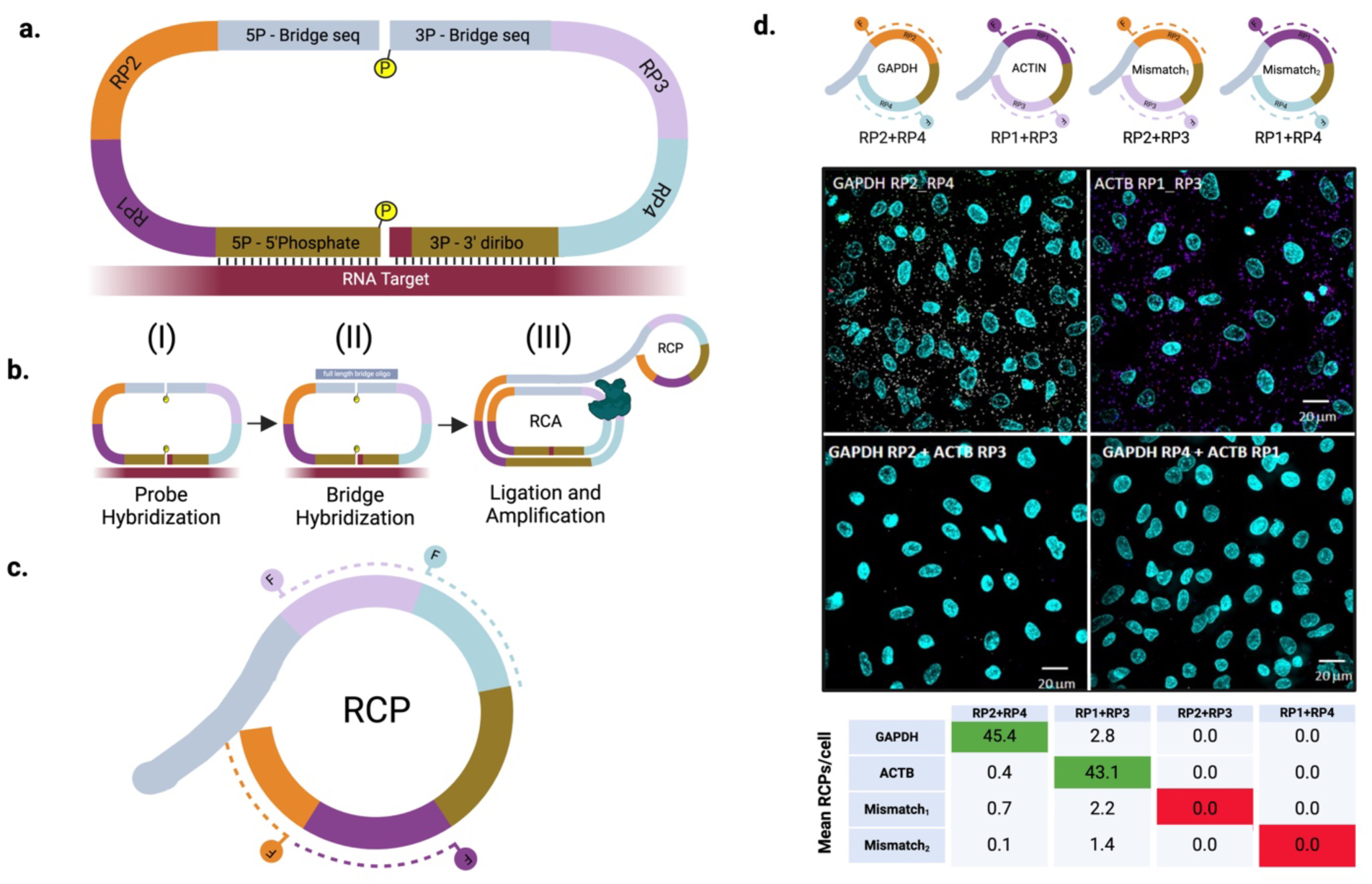
Bipartite LISH probes for spatial RNA analysis. **(a)** LISH-LnR design. 4 unique readout probe (RP) and universal 5P and 3P bridge sequences were appended to the 5P and 3P ends of LISH probes. (**b**) The LISH-LnR assay takes place in 3 steps. Ligation and amplification are combined in step III. (**c**) Each RCP is identified based on the fluorescence of 5’ fluorophores (circles, “F”) that are conjugated to complementary readout probe (RP) sequences (dotted). (**d**) LISH-LnR was performed on PFA-fixed A549 cells using a pool of probes targeting GAPDH and ACTIN with each probe set having a unique combination of RPs. Using a mix of fluorescently label oligos complementary to the individual RPs allowed detection and discrimination of both perfectly matched (GAPDH: RP2-RP4 and ACTIN (ACTB): RP1-RP3) and mismatched (Mismatch_1_: RP2-RP3 and Mismatch_2_: RP4-RP1) RP pairs. Fluorescent images were collected on the Olympus BX-51/Genus FISH Imaging System and decoded using the colocalization matched or mismatched RP sequences using Nikon-Elements AR. Image shows a single FOV representing all possible combinations of perfect and imperfect RP pair fluorescent signals. The number of overlapping signals was normalized to the cell number (number of DAPI puncta) per ROI. GAPDH and ACTB RCPs are pseudocolored in white or purple, respectively.

The LISH-LnR amplicon is prepared in three steps (**Fig. 1b**): (I) Probe sets are hybridized onto RNA targets and excess probes are washed away. (II) A universal probe set bridging oligonucleotide is hybridized to the 5P-Bridge and 3P-Bridge sequences. (III) The tissue sample containing the annealed probe sets and bridging oligonucleotide is incubated with a multi-enzyme reaction mix, consisting of two T4 ligases and the Phi29 DNA polymerase. The probe sets are circularized in a single reaction involving the combined action of the two ligases: T4 RNA Ligase 2 (Rnl2) and T4 DNA ligase, which seal the target RNA-templated junction and the bridging oligonucleotide-templated junction, respectively (thereby “locking” the circularized probe around the target RNA due to the twist of the hybrid helix). The Phi29 DNA polymerase uses the bridging oligonucleotide as a primer to initiate rolling circle amplification (RCA). The products of rolling circle amplification, which are produced proximally to their RNA target and become locally entangled in the fixed tissue, are subsequently available for detection via the many repeated copies of single-stranded readout probe binding sites that make up the rolling circle products (RCPs) (**Fig 1c**). The preparation of RCPs *in situ* can be conveniently completed in less than a day.

Low-plex imaging of RCPs can be performed on a standard fluorescence microscope. We used a 2-plex panel to assess colocalization of correctly matched versus mismatched probe pairs and their resulting RCPs. Single probe sets targeting the housekeeping genes *ACTIN* (3P-RP1 + 5P-RP3) and *GAPDH* (3P-RP2 + 5P-RP4) were used to assess this simplified version of the assay. The two probe sets were mixed and hybridized to paraformaldehyde (PFA)-fixed and permeabilized A549 cells. The number of colocalized fluorescent spots indicating perfect matches between correct RP pairs (*ACTIN*: RP1-RP3; *GAPDH*: RP2-RP4) and imperfect RP pairs (RP1-RP4 and RP2-RP3) were determined per cell. We detected 45.4 correctly colocalized *GAPDH* spots per cell and 43.1 correctly colocalized *ACTIN* spots per cell, with a median of 0 mismatched RCPs per cell (**Fig. 1d**). These data confirm the high specificity of matched probe pairing and illustrate how low-plex LISH-LnR assays can be analyzed using a standard microscope.

### Automation and tuning of high-plex LISH-LnR assays

High-plex LISH-LnR assays can be efficiently analyzed using combinatorial encoding schemas. Here, we incorporated sequential rounds of readout probe hybridization, imaging, probe stripping or photobleaching, and then finally image deconvolution with error correction. We used the Veranome Spatial Analyzer (VSA) for all high-plex assays in this study. The VSA performs sequential imaging and fluidic steps, then analyzes the combined images via a deep-learning algorithm trained to detect and decode individual RCPs (VeraWorks^TM^ 3.0) (**Fig. 2a-b**).

**Figure 2.**
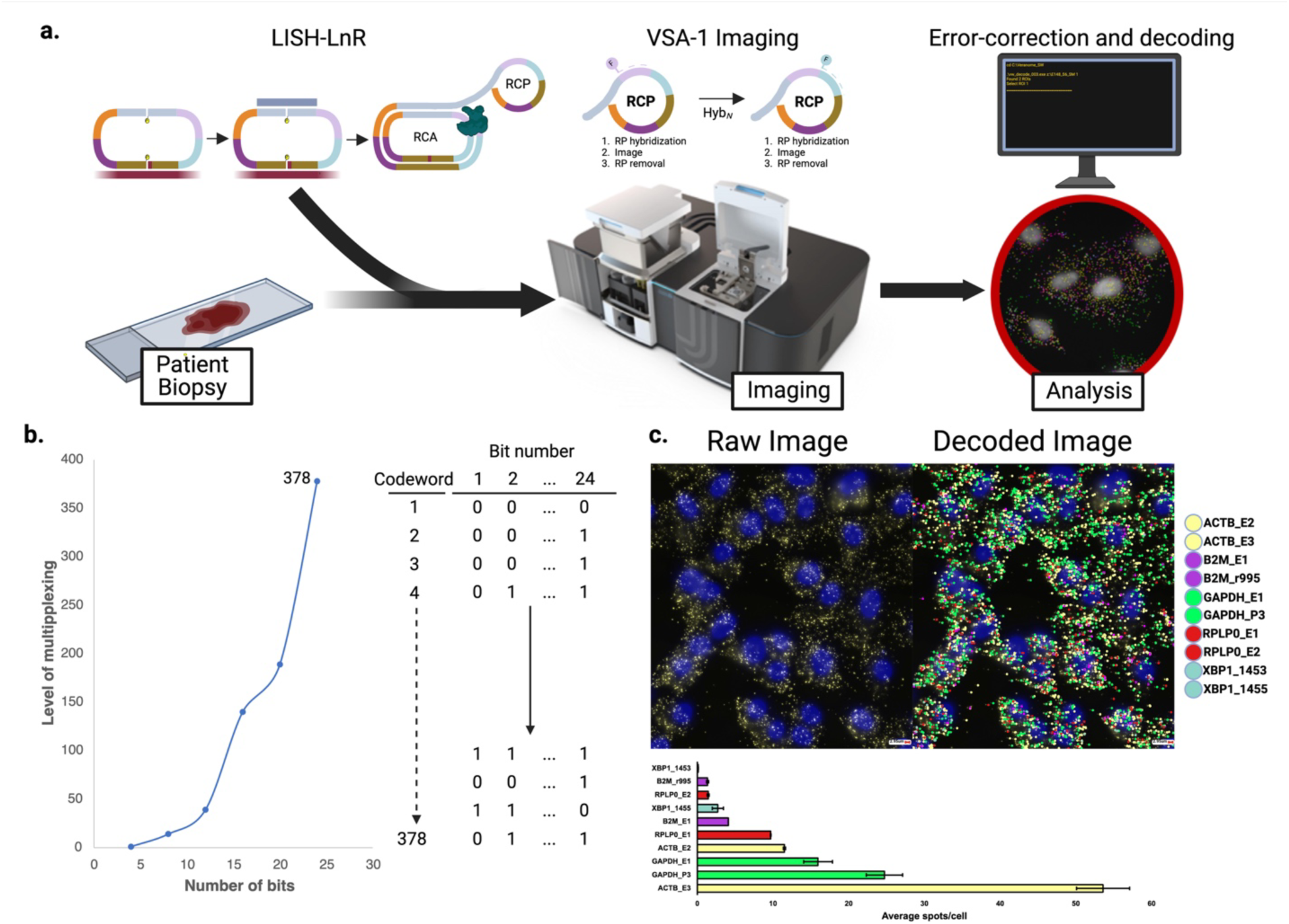
High-plex imaging of LISH-LnR products. **(a)** High-plex imaging pipeline. After samples are prepared using the LISH-LnR assay, they are placed on the VSA-1 (Veranome Biosystems LLC) where they undergo sequential rounds of RP hybridization, imaging, and RP removal. Images are then error-corrected, decoded, and analyzed **(b)** Codebook design. **(left)** Using a binary coding format (0/1) a code generation script was used to calculate the possible number of codewords (y-axis) for a given number of bits (x-axis). **(right)** Example of a 378-plex codebook with each codeword assigned to a 24 bit string of binary words. **(c)** A549 cells analyzed with the VSA-1 using a 10-plex housekeeping gene panel. Nuclei are labeled with DAPI (blue) and detected RNA molecules are indicated in yellow (raw image, left). RNA species were detected using VeraWorks 3.0 and individual RCPs labeled with pseudocolor overlays (decoded image, right).

The high-plex LISH-LnR probe sets used in this study were designed to carry two independent readout probe binding sites per probe (four independent sites per circularized probe set), thus enabling a 4-on bit set of codewords (**Fig. 1a**). Enforcing a minimum Hamming distance of 4,(*8*) we constructed a 24-bit codebook and assigned each mRNA target to one of 378 unique 24-bit codewords (**Fig. 2b**). To evaluate the detection and decoding of LISH-LnR based RCPs with the VSA, we first designed a house-keeping gene panel consisting of 10 probe sets targeting five genes (two probe sets per target). We detected ∼125 ±9 (Average ±SD) spots per A549 cell (**Fig. 2c**).

Anticipating physical constraints on the molecular size and number of RCPs in the setting of high-plex analyses, we investigated whether we could intentionally modulate these properties of the assay. We found that two strategies may be helpful to avoid molecular overcrowding. First, a probe decoy strategy may be employed to reduce the number of RCPs arising from specific abundant targets. In this strategy, a competing unligatable (“decoy”) 3P probe without the 3’-diribonucleotide is added to the hybridization reaction in a ratio desired to reduce RCP formation (**Fig. 3a**). We demonstrate the effectiveness of the decoy strategy by targeting the highly abundant 18S ribosomal RNA in A549 cells. We observed a specific decrease of 18S RCPs that roughly corresponds to the ratio of decoy over authentic probe used (**Fig. 3b**).

**Figure 3.**
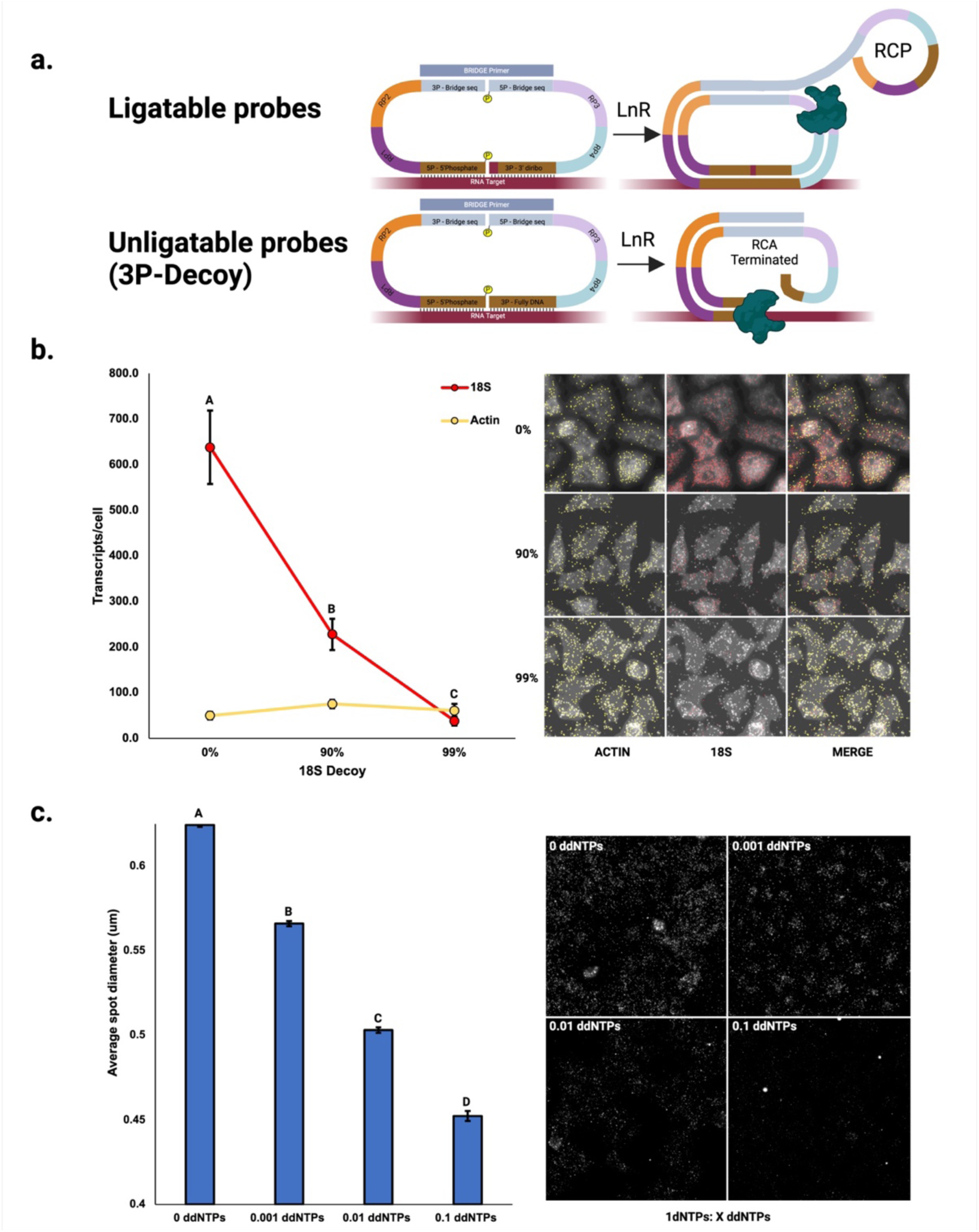
Molecular rheostats for tuning LISH-LnR. **(a)** Unligatable (decoy) 3P probes can be added to a hybridization reaction to selectively block RCP formation from the selected targets. **(b, left)** 90% and 99% 18S 3P decoy mix significantly reduce the number of 18S spots/cell [F(2,15)=72.9, p<0.001 Tukey post hoc HSD test]. The error bars represent SEMs. The number of ACTIN transcripts detected/cell is not affected by the 18S decoy probes [F(2,15)=2.8, p=0.09]. **(c)** A ratio of 1 dNTP: X ddNTPs was applied to A549 cells hybridized to a 16-plex immune-oncology panel. There is a significant effect of the dNTP: ddNTP ratio at the p<0.001 level [F(3, 40698)=1230, p<0.001]. Groups with different letter values (A-D) are significantly different as determined by the Tukey post hoc HSD test.

A second, independent approach to reduce potentially problematic molecular crowding is to globally reduce the size of the RCPs. This can be accomplished by inclusion of a chain-elongation inhibitors (chain terminator) such as ddNTP into the multi-enzyme reaction cocktail. Titration of ddNTP into the amplification reaction resulted in a dose-dependent decrease of spot size (**Fig. 3c**). Smaller spot size is expected to decrease deleterious effects associated with molecular crowding, but may reduce the sensitivity to detect smaller RCPs under the same imaging conditions. Decoying specific targets and/or globally reducing spot size therefore provides “molecular rheostats” that can be utilized to precisely tune LISH-LnR assay performance.

### Imaging and decoding high-plex LISH-LnR panels

We constructed a high-plex LISH-LnR immuno-oncology (I/O) panel, comprised of 372 probes targeting 130 frequently studied mRNA targets. As a first evaluation of the LISH-LnR I/O panel, we utilized the non-spatially resolved LISH-seq assay with FFPE tonsilitis samples. To this end, the bridging primer and T4 DNA ligase are excluded from the reaction, such that probe sets are ligated *in situ* only by Rnl2, creating linear, non-amplified ligation product. These linear ligation products are then released by RNase H digest of the probe-hybridized mRNA. PCR is used to amplify the released ligation products using universal primers, followed by analysis using DNA sequencing.(*6*) An average of ∼500,000 LISH-seq reads were obtained from each LISH-seq library where Rnl2 was included, but only 349 reads were obtained from a negative control sample that did not include the Rnl2 enzyme (-Rnl2). Since inexpensive qualification of LISH-LnR probe set designs can be of great utility, we assessed the concordance of less expensive, fully DNA probes ligated by SplintR versus 3’-diribo terminated probes for use with Rnl2. While fully DNA LISH probes ligated by SplintR can indeed serve as a proxy for Rnl2 LISH-seq, ligation occurs at a lower efficiency and with a lower correlation with bulk RNA-seq **(Fig. S1)**

Inclusion body myositis (IBM) is the most common inflammatory myopathy in patients older than 50. IBM is characterized by immune cell infiltration of the skeletal muscle and is typically diagnosed via biopsy. The nature of the immune infiltrate has not been characterized in detail. We performed LISH-seq on an IBM muscle biopsy specimen using our LISH-LnR I/O probe panel. LISH-seq data correlates strongly with bulk RNA-seq data (r = 0.866, p<0.001, **Fig. S2**). We next used the VSA to image LISH-LnR products generated *in situ* in fresh frozen (FF) samples (**Fig. 4 left, center**). This analysis revealed a very high expression level of IgG mRNA within the muscle tissue, consistent with a predominance of class-switched plasma cells, as has been previously reported.(*9*) Importantly, VSA image-based counting of the LISH-LnR products correlated well with both the LISH-seq results (**Fig. 4, right**, r = 0.917) and the bulk RNA-seq data (**Fig. S2**, r = 0.987).

**Figure 4.**
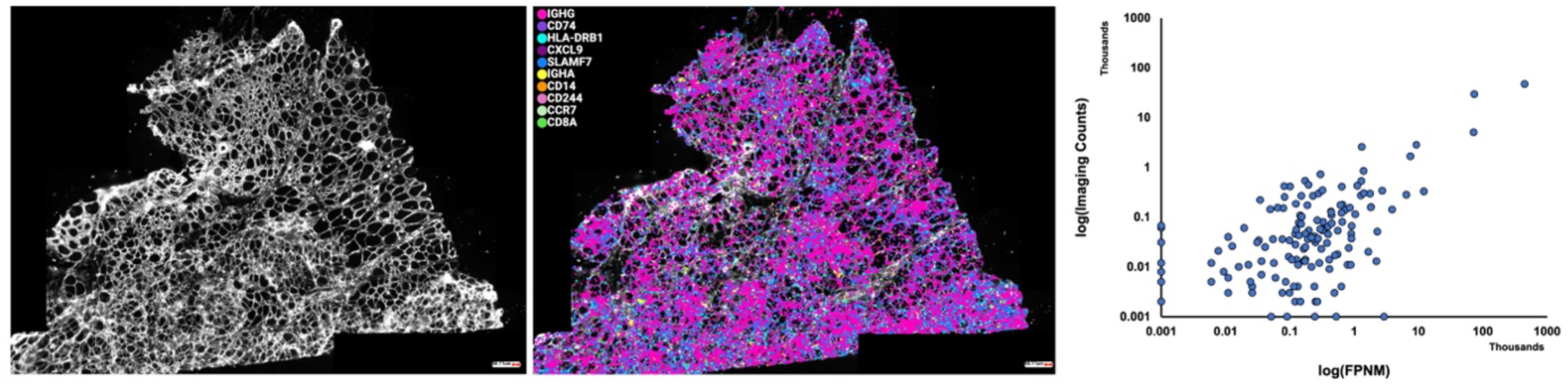
High-plex LISH-LnR in fresh fixed IBM samples. **(left)** IBM patient samples were stained with wheat germ agglutinin (WGA) to mark cellular membranes. **(center**) Pseudocolor overlay of the 10 most abundant LISH-LnR I/O RCPs detected: IGHG (4,594 transcripts/mm^2^), CD74 (2,892 transcripts/mm^2^), HLA-DRB1 (493 transcripts/mm^2^), CXCL9 (277 transcripts/mm^2^), SLAMF7 (252 transcripts/mm^2^), IGHA (162 transcripts/mm^2^), CD14 (83 transcripts/mm^2^), CD244 (71 transcripts/mm^2^), CCR7 (52 transcripts/mm^2^), CD8A (52 transcripts/mm^2^). (**right**) Serial sections of IBM fresh frozen muscle were prepared for LISH-LnR and for LISH-seq with the same probe panel design. FPNM was calculated from LISH-seq data (Methods) and was significantly correlated with LISH-LnR results (r = 0.917, p < 0.001).

### LISH-QC for rapid qualification of specimens before spatial transcriptomics

Vast archives of FFPE specimens are available at institutions globally. Current methods to assess RNA integrity, which rely on measuring bulk RNA, are poorly predictive of spatial transcriptomic assay success (*10*). This is especially problematic given the extreme pre-analytical variability of mRNA integrity across archival clinical specimens, coupled with the lack of standardized tissue-preparation protocols (*11*). Due to the high cost and low sample throughput of spatial transcriptomic assays, there is thus a critical need for a rapid, inexpensive, and reliable assay to pre-qualify FFPE tissue mRNA. Here we introduce LISH-QC, a simple and inexpensive qPCR-based assay to quantify FFPE mRNA abundance and accessibility for probe hybridization.

LISH-QC uses a housekeeping gene (here *ACTIN* or *GAPDH*) to template ligation of a fully DNA LISH probe set. Despite the lower efficiency of using SplintR and fully DNA LISH probes (**Fig. S1a**), this approach is compatible with direct qPCR quantification of the released ligation products using SYBR green incorporation with a standard Taq polymerase. (Ligation products derived from the more efficient Rnl2 ligation of diribonucleotide-containing LISH probes must alternatively be pre-amplified by a polymerase that is insensitive to inclusion of RNA-bases in the DNA template, eg. Herculase.) Using an optimized blend of SplintR ligase and RNAse H, LISH ligation products can be rapidly formed and enzymatically released for downstream real time qPCR quantification. The amount of released ligation product is then compared to a standard curve, and finally normalized to the surface area of the 10μm-thick tissue specimen on the slide (**Fig. 5a**). To assess the reliability of LISH-QC in qualifying accessible mRNA, we applied serial dilutions of RNase A (an endoribonuclease that degrades RNA at C and U residues) to FFPE tonsil sections for 10 minutes and then processed the samples using either LISH-QC, LISH-seq, or LISH-LnR. LISH-QC reported a linear decrease in signal from both *GAPDH* and *ACTIN* probe sets with increasing RNase A concentration (**Fig. 5a**). In this experiment, LISH-QC data was therefore predictive of LISH-LnR and LISH-seq assay performance (**Figs. 5b, 5c** respectively).

**Figure 5.**
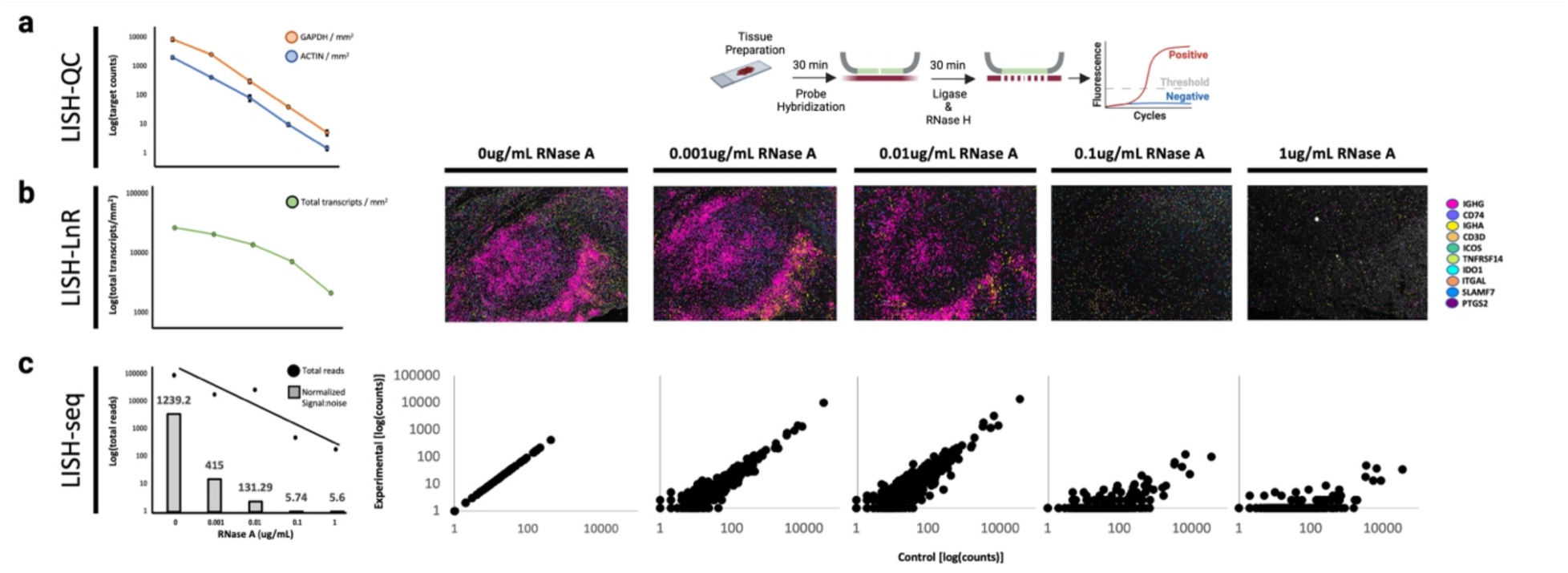
LISH-QC qualifies FFPE RNA for transcriptomics assays. **(a, right)** Schema of the LISH-QC assay. After RNA is made accessible, fully DNA ACTIN and GAPDH probesets are hybridized onto biological samples for 30mins. Probes are then simultaneously ligated and removed in a SplintR-RNase A reaction mix. Eluted probes are then quantified using qPCR. **(a, left)** Triplicate FFPE tonsilitis samples were incubated in 0, 0.001, 0.01, 0.01, or 1ug/mL RNase A for 10 minutes at 37°C. RNase A reduced GAPDH [F(4,10)=73.4, p<0.001] and ACTIN [F(4,10)=27.2, p<0.001] target molecules. **(b)** Representative images are shown to the right, displaying the most abundant transcripts in control FFPE tonsilitis. Imaging counts are highly correlated to LISH-QC quantified GAPDH (r = 0.806, p<0.001)) and ACTIN (r = 0.826,p<0.001)) molecules over the same dilution series. **(c)** The overlayed bar graph indicates the number of matched:mismatched sequencing reads for each RNase A treatment. Read counts are highly correlated to LISH-QC quantified GAPDH (r = 0.949, p<0.001) and ACTIN (r = 0.905, p<0.001)) molecules. Plots to the right display the correlation of LISH-seq data between the control (0ug/mL) total read counts and the corresponding RNase A treatment.

### Spatial mapping of mRNA splice variants with LISH-LnR

Alternative splicing events lead to the expression of diverse mRNA isoforms and the translation of different protein products. There remains an unmet need for a technique to spatially localize mRNA isoforms of interest (*12–16*). Because single LISH-LnR probe sets are capable of producing detectable signals, and their specificity can be readily designed to discriminate between different exon junctions, we investigated the utility of LISH-LnR for mRNA isoform localization in FFPE tissues. As a proof-of-concept, we designed probe sets to detect two alternative isoforms of X-box binding protein 1 (*XBP1*), which is a key modulator of the unfolded protein response (UPR). The unspliced variant of *XBP1* (XBP1-unspliced, “XBP1-u”) contains 7 exons (*17*) and is the dominant isoform in cells with low amount of endoplasmic reticulum (ER) stress. In states of high ER stress (hypoxia, nutrient deficiency, excessive ROS, decreased ATP, proteostatic stress, etc.), XBP1-u undergoes unconventional splicing, in which a small intronic 26 nucleotide sequence from exon 4 is removed by the endoribonuclease, IRE1, to create “XBP1-s,” which contains a translational frameshift that leads to the expression of functional protein. We designed 3 LISH-LnR probes to detect the presence and location of both splice variants (**Fig. 6a**). Previous studies have demonstrated that XBP1 splicing can be induced by thapsigargin, which evokes ER stress by inhibiting the ER Ca^2+^ ATPase (*18*). After treatment of A549 cells with 50nM of thapsigargin (Tg), the total number of XBP1 spots per cell did not change, but the ratio of XBP1-s to XBP1-u increased significantly (**Fig. 6b)**.

**Figure 6.**
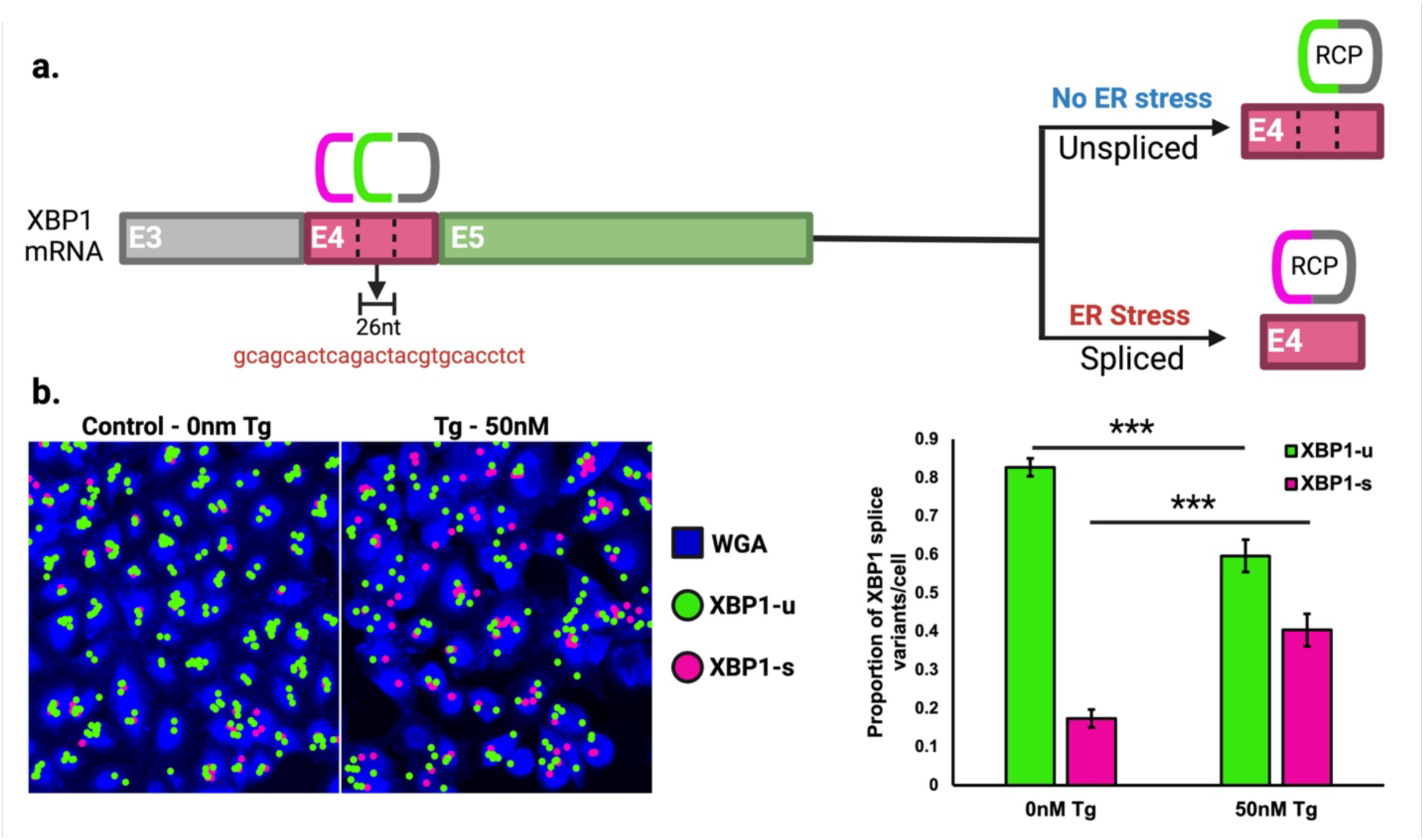
Spatial analysis of XBP1 splicing with LISH-LnR. **(a)** 3 LISH-LnR probes were designed to detect both XBP1 isoforms: 3P ligation probes recognize either the spliced (magenta) or unspliced (green) variant. When ligated to the shared 5P ligation probe (grey) and amplified during the LISH-LnR assay, unspliced and spliced rolling circle products have distinguishable readout probe combinations. **(b)** A549 cells were incubated with 50 nm thapsigargin or vehicle control (DMSO) for 3 hours and then assayed for XBP1 splicing using LISH-LnR probes. Fluorescent images are shown to the left, quantified to the right. The proportion of spliced XBP1 transcripts increased with the thapsigargin treatment (*t*(138)=5.04, *p*<0.0001).

Having validated the XBP-1 LISH-LnR assay in fixed and permeabilized cells, we next sought to localize XBP1 isoforms in FFPE patient samples. Rhabdomyosarcoma (RMS) is the most prevalent form of malignant soft tissue sarcoma in children and is associated with poor survival outcomes (*19*). It is well documented that RMS cells activate the ER stress inducers IRE1 and PERK, as well as their downstream signaling elements ATF4 and XBP1-s (*20*). Within the RMS tumor bed, tertiary lymphoid structures (TLS) are frequently observed, but the relationship between immune cell activation and RMS ER stress has yet to be elucidated. We therefore identified archival FFPE specimens from two RMS patients that each contained definitive TLS sites. The RNA quality of these blocks was assessed by LISH-QC and found to be acceptable for transcriptomic assays (**Fig. 7a**). LISH-Seq and LISH-LnR analysis using the I/O panel, supplemented with the XBP1 transcript isoform probes, yielded a Pearson correlation coefficient of R ∼ 0.85 for both sites (**Fig. 7b-c**). Further, LISH-LnR spot counts revealed a strong correlation between the two RMS specimens (**Fig. 7d**). Unsupervised Leiden clustering (*21*) of the LISH-LnR data detected discrete immune cell populations, which were autocorrelated in both blocks (**Fig. 7e, Fig. S3**).

**Figure 7.**
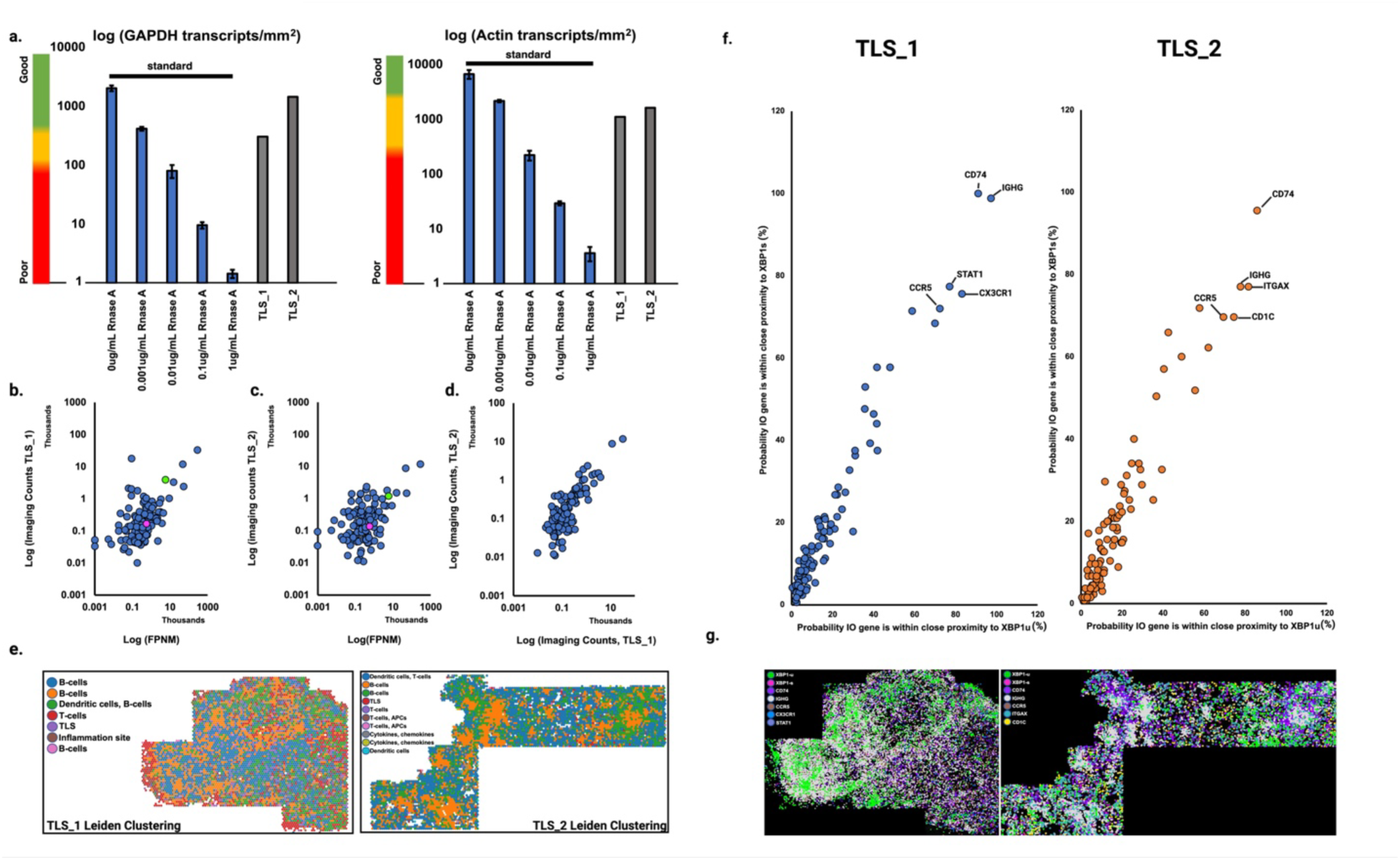
Spatial transcriptomic analysis of RMS. **(a)** LISH-QC was performed on 2 FFPE RMS patient blocks containing tertiary lymphoid structures (TLS_1 and TLS_2). **(b, c)** There is a significant positive relationship between the two TLS patient blocks (TLS_1, TLS_2) and LISH-seq data prepared from a single TLS site. TLS_1: r = 0.86, *p* < 0.001; TLS_2: r = 0.845, *p* < 0.001). The location of each XBP1 isoform is indicated in each plot, where XBP1-s is shown in magenta and XBP1-u is shown in green. **(d)** There is a strong correlation between imaging counts of TLS_1 and TLS_2 (r = 0.937, *p* < 0.001). **(e)** Transcriptomics data was binned into 50 μm^2^ grids and spatially unaware analysis (Leiden clustering) followed by Wilcox Rank sum tests were used to evaluate the transcriptomics profile of each bin. Cell types associated with each cluster (annotated) were assigned based on *a prior* knowledge of typical cell transcriptomics profiles infiltrating oncology tissue. 7 clusters were identified in TLS_1 and 10 clusters were identified in TLS_2. (**f**) Probability (%) that an I/O gene appears within 60 um of a XBP1s (y axes) or XBP1u transcript (x axes). (**g**) Pseudocolor overlays of genes that are likely to appear within close proximity to an XBP1 variant.

Changes in gene expression associated with XBP1 splicing may be observed by determining whether an I/O gene is more likely to fall in close proximity (within 60 μm) to XBP1s versus XBP1u. To evaluate this, we calculated the distance of the closest transcript (‘near neighbor’) for each member of the LISH I/O panel to each XBP1u and XBP1s transcript identified in TLS_1 and TLS_2 and then determined the probability each I/O gene falls within close proximity to the variant. As shown in **Fig. 7f**, there are no major outliers to suggest significant changes in I/O gene expression associated with XBP1 splicing; for both TLS_1 and TLS_2, the probability that an I/O gene will segregate in close proximity to XBP1s and XBP1u is roughly equivalent. I/O genes that are likely to segregate in close proximity to XBP1s and XBP1u (>70% probability) are annotated in the LISH-LnR imaged regions (**Fig. 7g**). While we do not observe focused regions of XBP1-s, we do note that XBP1-u expression appears elevated and colocalize within the TLSs themselves, which contains abundant B cells (based on class-switched immunoglobulin, IGHG), antigen presenting cells (based on CD74), and T cells (based on CX3CR1, CCR5) (**Fig. 7g, Fig. S4)**.

## DISCUSSION

Here we describe a novel Ligation *In Situ* Hybridization (LISH) assay that uses circularization and rolling circle amplification to locally produce many copies (100’s to 1000’s) of readout probe binding sites, which can be visualized via combinatorial hybridization with fluorescently tagged oligonucleotide probes. We demonstrate how low-plex LISH-LnR panels can produce detectable and decodable spots using standard microscopy and analytical software. LISH-LnR is also amenable to analysis of high-plex panels using automated fluidics and imaging instrumentation. We used the Veranome Spatial Analyzer and its integrated software to perform spatial transcriptomic analyses of cells, fresh-frozen tissues, and archival FFPE specimens.

The LISH-LnR spatial assay is augmented by integration with two related assays: LISH-QC and LISH-seq. LISH-QC, introduced here, is a simple, rapid, low-cost assay to quantify the number of available housekeeping mRNA target sites within a sample. LISH-QC addresses a critical need for sample qualification prior to running high complexity, high cost, low throughput spatial transcriptomic assays. It is particularly critical to qualify archival FFPE specimens, which typically suffer from highly variable pre-analytical fixation and storage effects that impact RNA quality. Notably, the LISH-QC assay qualifies RNA availability at the stage just prior to high-plex analysis. Therefore, both RNA quality and tissue pre-treatment protocols are tested, providing information that is highly predictive of downstream hybridization-based assay performance. Finally, while not demonstrated here, the LISH-QC assay can in theory be performed on the same tissue section that is used for spatial transcriptomics, with the caveat that the housekeeping gene target sites will no longer be available and the RNAse H must be thoroughly washed away prior to the hybridization of a probe panel. Integration of LISH-seq with spatial LISH-LnR assays can also provide valuable information in at least two ways. First, LISH-seq can be used to pre-qualify probe designs using inexpensive surrogate probe sets. These surrogates need not incorporate the readout probe binding sites, since sequencing will be used as a readout. Further, they need not include the 3’ diribonucleotide modification since the SplintR ligase can serve as a less efficient proxy. The design of the surrogate probe sets can be tested via deep sequencing to determine their matched versus mismatched ligation rates, and to compare their performance against available RNAseq data. Second, LISH-seq data can be used to serve as ground truth for the optimization of LISH-LnR image analysis. For example, discordance with LISH-seq data may indicate a problematic optical artifact associated with deconvolution of high plex fluorescence data.

A unique advantage of the LISH-LnR assay is the flexibility to incorporate “molecular rheostat” control over RCP number and size. We demonstrate that including a ddNTP along with Phi29 polymerase can reliably reduce the size of the RCPs. It is important to note, though, that the theoretical advantage of packing in more spots per unit volume might be traded off a loss of signal-over-background, which may limit the rate at which a given tissue area can be imaged, for example. We also demonstrate the use of fully DNA acceptor probes to reduce, via competition with diribo-containing acceptor probes, the number of target-specific RCPs. Other forms of decoy probes could also be developed, including donor or acceptor probes that lack their 5’-phosphorylation. The RNA target site could also be blocked using an oligo, but this approach might exhibit less linear behavior. While not explored here, we expect that the use of multiple probe sets per target mRNA will additively increase RCPs per target transcript.

mRNA splicing contributes enormous diversity to eukaryotic transcriptomes and proteomes. While splicing events occur naturally in cells, misregulation of gene splicing has been linked to various diseases including cancer (*22*), neurodegeneration (*23, 24*) and myopathy (*25*). Spatial localization of splice variation is an important objective for *in situ* hybridization platforms, but most approaches are not amenable to detecting inclusion or exclusion of shorter sequences (*12–16*)). By virtue of their high signal over background, we see that individual LISH-LnR probe sets can be used to detect biologically important splicing of a 26nt sequence from the XBP1 mRNA. After validating our probe design in cell culture, we produce high-plex spatial LISH-LnR data sets from RMS patient FFPE specimens containing tertiary lymphoid structures. We observe high levels of unspliced XBP1 mRNA localized to the TLSs, which is not accompanied by large amounts of spliced XBP1. This observation is consistent with a scenario in which XBP1 is expressed at high levels in the TLS, where it is not constitutively spliced, but is available for a rapid shift to the spliced form should ER stress rise.

In summary, recent advances in spatially resolved transcriptomics have expanded our knowledge of complex tissue architectures in health and disease. The ecosystem of LISH assays introduced here comprises a powerful platform to advance spatial biology. With probe set specificity capable of distinguishing transcript isoforms and the flexibility to tune signal strengths up or down, we expect that LISH-LnR assays will ultimately find utility in highly optimized clinical diagnostics.

## METHODS

### Imaging and analysis of low-plex LISH-LnR

LISH-LnR low plex panels (2-plex) were imaged on an Olympus BX-51/Genus FISH Imaging System using filters and exposure times appropriate for the fluorophores hybridized to the given readout probes (sequences and conjugated fluorophores are listed in **Table S1**). Imaging data was transferred to Nikon-Elements AR, where the codewords of all probesets (and mismatched words if relevant) were implemented into the detection of each RCP species. RCPs were decoded based on the colocalization of probe set-specific RP-conjugated fluorophores (2).

### Imaging and analysis of high-plex LISH-LnR

The VSA-1 (Veranome Spatial Analyzer) was configured and prepped according to the manufacturer’s instructions with manufacturer supplied reagents (200ml Cycling Buffer, Hybridization Buffer, Imaging Buffer A and B, Wash Buffer 1 and Quenching Buffer 2) and readout probes. Readout probes were added to 5 mL of Hybridization Buffer for a final readout probe concentration of 20 nM. All fluidics were controlled and calibrated according to the manufacturer. Sectioned samples on 40 mm poly-L-lysine coated slides were removed from storage at 4°C and loaded into a flow cell with 300 μL of 2X-SCC. The flow cell was loaded into the VSA-1 imaging chamber, protected from light exposure. An initial scan of the samples was performed using 640 nm emission at 10% power, scanning the center of the sectioned tissue thickness. Two regions of interested were selected from the initial scan to be imaged across 9 vertical steps of 1 um each, resulting in an imaging depth of 8 um. Readout probes were hybridized on the sample within the imaging chamber for 25 minutes at room temperature, followed by imaging with the following laser excitation: 750 nm at 75% power for 125 ms, 640 nm at 20% power for 100 ms, 577 nm at 30% power for 100 ms, and 530 nm at 50% power for 100 ms. The readout probes were removed using chemical quenching with Quenching Buffer 2 (following the manufacturer’s instructions), allowing fresh readout-probes to be hybridized for the next imaging round. 7 rounds with 24 unique readout-probes and 2 landmark probes (LMPs) were completed. DAPI (0.5mg/mL in 1x-PBS) and wheat germ agglutinate (WGA, 1 µg/mL in 1x-PBS) were incubated on the sample for 15 minutes at room temperature and imaged using 470 nm at 5% power for 50 ms and 640 nm at 5% power for 50 ms. Analysis (spot registration and decoding): image analysis was performed with Verascape software with the following system requirements. Intel Xeon Silver 42124R processor, NVIDIA RTX A4500 graphic card, Python 3.8, and MATLAB runtime R2019a V9.6. Decoding software was run following commands developed by the manufacturer. Analysis (Veraworks Version 3): RNA transcript codewords and 10% negative control codewords following minimum Hamming distance 4 (MHD4) across 24 unique readout probes were created. Spots were registered and decoded individually (“slice-by-slice”) across 4µm. Analysis (Spatial analysis): Decoded RNA transcripts are analyzed further using Verascape with 25% MisID. Transcript profiles were visually QCed and obvious false positive genes were removed from the analysis.

### Cell lines and tissue specimens

A549 lung epithelial cells (catalog# CRM-CCL-185) were purchased from ATCC (Manassas, VA). A549 cells were cultured in DMEM (Thermo Fisher Scientific) supplemented with 10% FBS (Thermo Fisher Scientific), and glutamine (Thermo Fisher Scientific) and Penn/Strep (Thermo Fisher Scientific) on circular glass coverslips coated with poly-D-lysine (Millipore) for 24 - 48 hours at 37°C with 5% CO_2_ in a humidified incubator. Human tonsil FFPE blocks (catalog# AMS6022) were purchased from ASMBio (Cambridge, MA). Tissue blocks were stored at -20C°. The use of muscle biopsy samples (IBM and control) was approved by the Johns Hopkins Institutional Review Board (IRB00072691). FFPE tumor tissue blocks were obtained from the pathology archives at Johns Hopkins Hospital (Baltimore, MD) from patients diagnosed with alveolar rhabdomyosarcoma. The tissue collection protocol was approved by the Johns Hopkins Institutional Review Board (IRB).

### Ligation probe and RP sequences

All ligation probe and RP sequences used for this paper can be found in **Table S1.** Ligation probe sequences were identified using www.primer3plus.com parameters in Supplemental File <u>primer3settings.txt</u>.

### LISH pretreatment protocols

All the following preparation protocols are performed immediately before the i. LISH-LnR, ii. LISH-QC, or iii. LISH-seq assays. Cell cultures: 8×10^4^ A549 cells (ATCC, ccl-185) were plated onto 40mm glass cover slips, fixed with 4% PFA, and frozen in -80°C overnight. The next morning, cells were washed 3x in 1x-PBS (100 μL/wash) and permeabilized in 100 μL perm buffer [0.1% Triton X-100, 1x PBS, 100 U/mL RNase Protector (Sigma-Aldrich)]. After 20 minutes, cells were incubated in 0.1 N HCl for 10 minutes. 100 μL of 1x PBS was then used to wash the cells a total of 3x.

Formalin-Fixed Paraffin-Embedded Tissues (FFPE) Pretreatment: 4 μm or 10 μm thick sectioned FFPE tissue samples were removed from -80°C and immediately transferred to a slide rack. Slides were baked in a dehumidified oven at 60°C for 1 hour. After the heated incubation, the slide rack was placed in a tissue-tek container filled with Xylene. The slide rack was allowed to incubate for 5 minutes, after which it was gently lifted and placed in a fresh container of Xylene. After a second 5-minute incubation, the slide rack was removed and placed in 100% ethanol for 1 minute. The slide rack was then transferred to a fresh container of 100% ethanol and incubated for 1 minute. The slide rack was then removed and set on a laboratory bench to air dry for 5 minutes at room temperature. The slide rack was then moved to a container with boiling, distilled water. After 10 seconds, the slide rack was moved to a container containing boiling 1x sodium citrate H.I.E.R (BioLegend, 928502). After 10 minutes the slide rack was transferred to a separate rinse container with distilled water. After the slides rinsed for 15 seconds, the slide rack was transferred to 100% ethanol for 3 minutes and then dried in a 60°C incubator for 5 minutes. Once dry, slides were removed from the rack and a pap pen (Abcam ab2601) was used to trace around the tissue region of interest. If prepared for LISH-QC, pictures of the tissues were captured. After a hydrophobic barrier was formed with the pap pen application, 2 μg/mL Proteinase K was applied to each section, completely covering the tissue. Samples were then incubated at 45°C. After 15 minutes, the protease was aspirated off and the tissues were immediately washed 3x with 1x PBS (0.5 mL/sample). Samples were then fixed with 4% PFA (200 μL/sample) for 5 minutes at room temperature. PFA was removed with 3 rinses of 1x PBS (0.5 mL/wash). Fresh Frozen (FF) Tissues: Sections were removed from -80°C and placed in a slide rack. The slide rack was then immediately transferred to container containing chilled (4°C) 10% formalin. After 15 minutes, the slide rack was transferred to 50% ethanol. After 5 minutes, the slide rack was then transferred to 70% ethanol. After a 5-minute incubation, the slide rack was transferred to 100% ethanol. After 5 minutes, the slide rack was transferred to a fresh container of 100% ethanol for 5 minutes. After the ethanol dehydration steps, slides are dried for 5 minutes at room temperature. Once dry, they are removed and tissues are outlined using a pap pen (Abcam ab2601).

### LISH-LnR

50 μL of ligation probe reaction [100 nM each 3P and 5P probe, 20% Formamide, 2x SSC, 20U Protector RNase Inhibitor (Roche) in UP H_2_0] is added to each tissue specimen. Coverslips are then applied to the samples, which are placed in a humidified chamber at 45°C. After a 4-hour incubation, coverslips are removed and the ligation probe reaction is washed twice with preheated (45°C) post ligation wash buffer (0.5 mL/wash: 20% Formamide, 2x SSC). Tissue specimens are then washed twice in 0.1x SSC (0.5 mL/wash/sample) and 50 μL full length bridge ligation mix (full length bridge primer suspended in 1x PBS at a concentration of 100 nm multiplied by the plex incorporated in the ligation probe reaction) is applied. Coverslips are used to cover the samples, which are then placed in a 37°C humidified chamber for 1 hour. After the bridge incubation, samples are washed in 0.1x SSC (0.5 mL/sample) followed by a 1x Phi29 buffer (Lucigen, 30221-2) exchange using 100 μL/sample. Phi29 + T4 Ligase + Rnl2 triple enzyme cocktail [1x Phi29 buffer (Lucigen, 30221-2), 800U T4 DNA Ligase (NEB M0202S), 300U Rnl2 (Enzymatics, L6080L), 1 mM ATP (NEB), 1 mM dNTPs (Applied Biosystems, 362271), 40U Protector Rnase Inhibitor (Sigma-Aldrich), 200U Phi29 (Lucigen 30221-2)] is prepared in a 100 μL volume and 50 μL is applied to each sample. Coverslips are then applied and samples are left to incubate overnight in a 30°C humidified chamber. The next morning, coverslips are removed and samples are washed three times in 2x SSC (0.5 mL/sample). Tissues are stored in 3 mL 2x SSC prior to imaging.

### LISH-QC

Hybridization, ligation and release of LISH-QC probes: 50 μL LnR Probe Hybridization mix (100nM each Actin and GAPDH 3P and 5P ligation probes in 20% formamide, 2x SSC) is applied to FFPE tissue samples prepared as described above. Samples are then placed in a humidified chamber at 37°C. After a 30-minute incubation, samples are removed and washed 3x with 100 μL preheated (37°C) 1x SplintR buffer (1x SplintR Ligase Buffer, NEB B0375S). 100 μL SplintR Reaction Buffer [1x SplintR Ligase Buffer (NEB), 1000U SplintR (NEB), 0.25U RNase H (NEB)] was then added to the samples at 37°C. After 30 minutes, the supernatant was collected. <u>qPCR:</u> LISH-QC products are analyzed on the QuantStudio 6 Flex (Thermo Fisher Scientific) using Brilliant III Ultra-Fast Sybr (Agilent, cat # 600882) mastermix. Standard curves are prepared using 8 dilutions (10^1^ to 10^8^ molecules/μL) of synthetic GAPDH and ACTIN, respectively. The concentration of ligated GAPDH and Actin molecules were determined based on their respective standard curves. Final concentrations were normalized to the tissue area so that the number of GAPDH and Actin molecules/mm^2^ was reported for each sample. GAPDH and Actin ligation, synthetic, and primer sequences used for qPCR can be found in **Table S1**.

### LISH-seq

Methods for LISH-seq follow the same workflow as LISH-LnR with the following exceptions. Following the three washes in 0.1x SSC (‘LISH-LnR assay’), samples are washed in 1x RnL2 buffer (100 μL/sample) and incubated in 50 μL RnL2 reaction buffer mixture [1x RnL2 Buffer, 150U Rnl2 (Enzymatics, L6080L), 20U Protector RNase Inhibitor (Roche)] overnight at 37°C. The following morning, samples are washed 2x in 2x SSC (0.5mL/sample) followed by 100 μL 1x RNase H buffer (NEB). 50 μL RNase H reaction mixture is then applied (1x RNase H buffer, 0.25U RNase H) to each sample and allowed to incubate at 37°C. After 30 minutes, the eluate is collected and stored at - 80°C until sequencing. Illumina Sequencing analysis: FASTQs were aligned to a file containing sequences of ‘perfect match’ and ‘mismatch’ ligation products, the latter which forms when the 3P and 5P probes of different gene targets ligate. Reads for all perfect matches and mismatches were tallied. FPNM calculation: Total reads in a sample (perfect matches and mismatches) were summed and then divided by 1,000,000 (a ‘per million’ scaling factor). Each read count was divided by the per million scaling factor (‘FM’), which was then divided by number of probes used to target the given gene (‘FPNM’).

### Statistics

Information regarding the quantification and statistical analyses performed on all data are described this section or the figure legends. A Person’s correlation coefficient (R) was used to measure the linear correlation between x and y variables (**Figs. 4, 5, 7; S1, S2).** A 2-way repeated measures ANOVA followed by a Tukey post hoc HSD test was incorporated in **Figs. 3, 5** and a paired-samples t-test was used to evaluate the data for **Fig. 6**. Genes specific to each cluster in **Fig. S3** were identified by Lieden were found using a Wilcoxon Rank sum test: (https://scanpy.readthedocs.io/en/stable/generated/scanpy.tl.rank_genes_groups.html, https://www.sc-best-practices.org/cellular_structure/annotation.html). <u>Leiden Clustering:</u> Spatial transcriptomics data was binned into 50um^2^ grids and Leiden Clustering (*21*) was used to evaluate the transcriptomics profile of each bin (Squidpy).

## ACKNOWLEDGEMENTS

We would like to thank our collaborators at Veranome Biosystems: Serdar Tulu, Marc Glazer, Chloe Kim, Shauna Edridge, Hareem Maune, Brain Hilbush, Bongjun Son. George McNamara (Ross Fluorescence Imaging Core) provided critical support on the Olympus BX-51/Genus FISH Imaging System. Oncology Tissue Services (OTS) at JHMI was a critical resource for our FFPE sample preparation. In particular, we would like to thank Wanda Stirling for sectioning all of our samples.

---

**Supplementary Figure 1.**
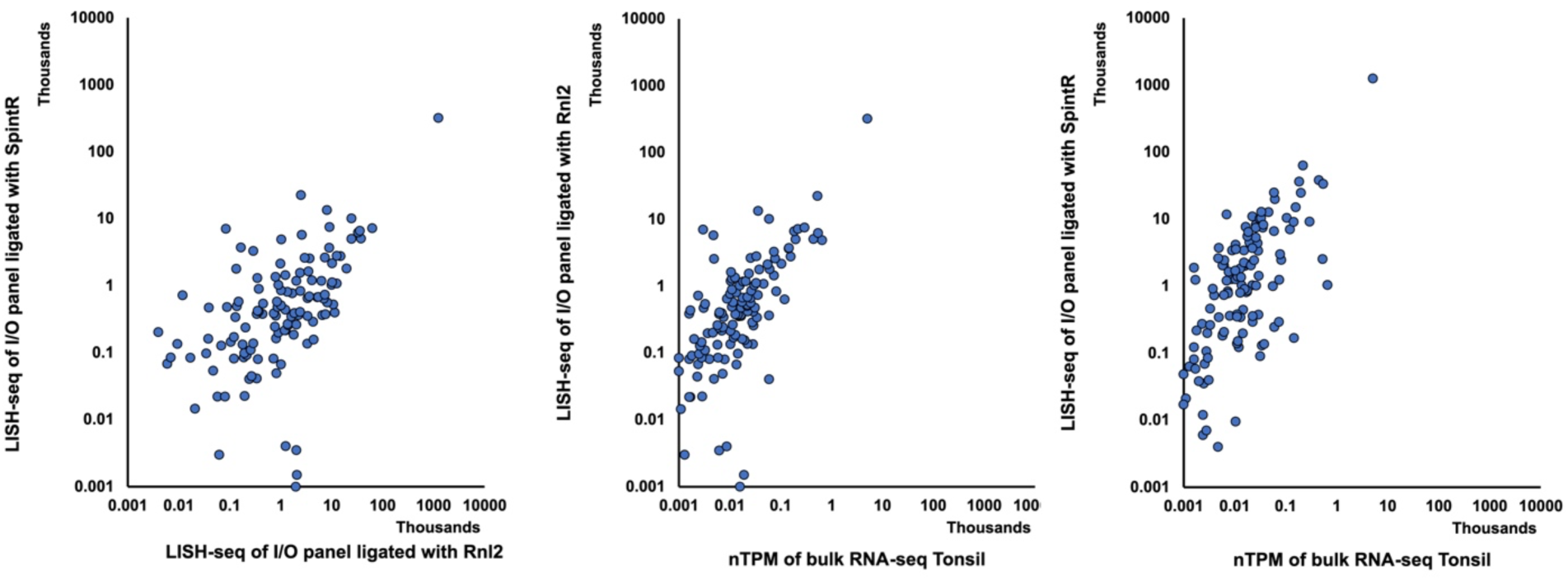
**(left)** Correlation between the average read count for 122 ligated probes generated from assays in which the same ligation probe sequences were ligated with SplintR (vertical axis - probes lack the diribonucleotide) or Rnl2 (x-axis - probes contain the diribonucleotide). 4 FFPE samples from tonsilitis were used for this experiment (2 samples for SplintR and 2 for RnL2). The read count for each ligated probe was averaged between the two samples. There is a significant correlation between the probesets ligated with SplintR and RnL2, r = 0.99, p<0.001. While the high correlation (r = 0.995) between the assays indicates that SplintR can be used as a proxy to evaluate ligation probe binding efficiencies prior to ordering as part of a LISH-LnR panel, Rnl2 LISH-LnR probe sets (**middle**) show slightly better correlation with bulk RNA-seq tonsil data (r = 0.985, p<0.001)) than fully DNA probe sets ligated by SplintR ((r = 0.981, p<0.001) (**right**).

**Supplementary Figure 2.**
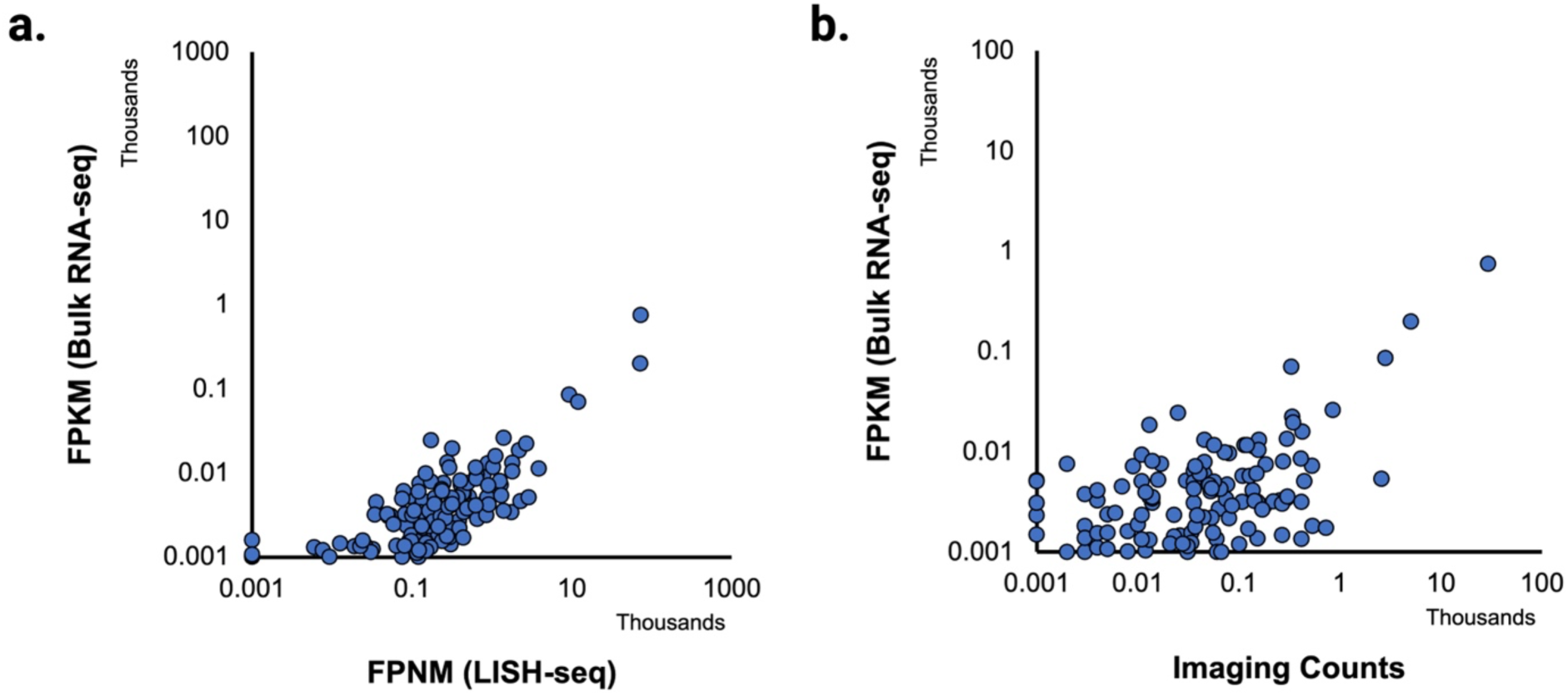
**(a)** Bulk RNA-seq was prepared from IBM patient samples. The LISH-LnR I/O panel was hybridized and ligated *in situ.* Ligated probes were released and sequenced. FPNM (Methods) is a modified calculation of FPKM where read counts are additionally normalized to the number of probes/gene. Bulk RNA-seq and LISH-seq data are correlated (r = 0.866, *p* < 0.001). **(b)** LISH-LnR imaging counts are correlated with bulk RNA-seq (r = 0.987, *p* < 0.001).

**Supplementary Figure 3.**
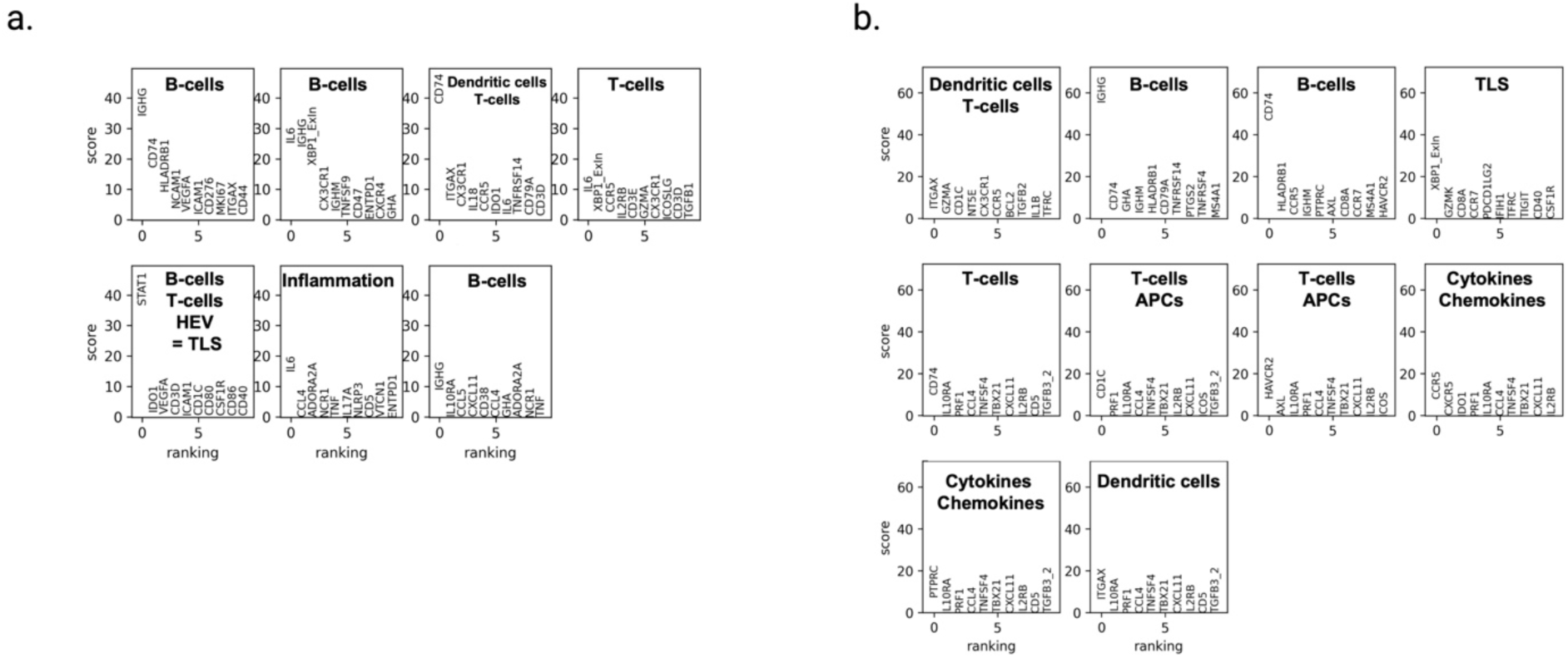
Results of the Wilcox rank tests from Leiden clustering **(a)** TLS_1 **(b**) TLS_2. Abbreviations for identified cell populations: Antigen Presenting Cells (APCs), High Endothelial Venules (HEV). Genes specific to each cluster were found using a Wilcox Rank sum test. Scores refer to ‘z-scores’ such that the higher the score, the more dominant the gene is within the cluster. 10 genes are annotated for each cluster. The cell type(s) representing each cluster were identified based on typical cellular transcriptomics signatures enhanced in oncology tissue (*26–32*). Both patients show clear immune activity: the presence of B-cell, T-cell types with associated inflammatory molecules, cytokines, and chemokine and chemokine receptors indicate active immune responses.

**Supplementary Figure 4.**
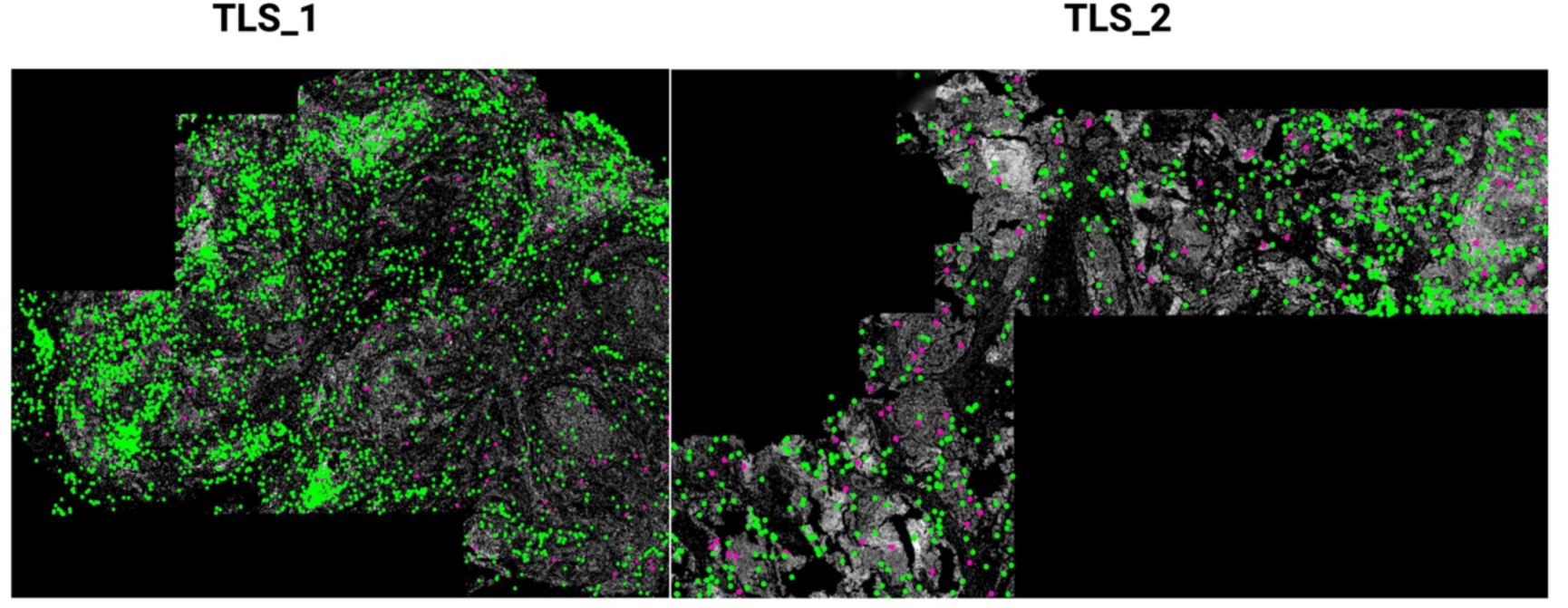
XBP1-u (green) and XBP1-s (magenta) overlays of TLS_1 (left) and TLS_2 (right). Cells in each region are stained with DAPI (shown in white).

